# Identification of a terpene synthase arsenal using long-read sequencing and genome assembly of *Aspergillus wentii*

**DOI:** 10.1101/2024.09.23.614465

**Authors:** Richard Olumakaiye, Christophe Corre, Fabrizio Alberti

## Abstract

Fungi are talented producers of secondary metabolites with applications in the pharmaceutical and agrochemical sectors. *Aspergillus wentii* CBS 141173 has gathered research interest due to its ability to produce high-value norditerpenoid compounds, including anticancer molecules. In this study, we aimed to expand the genomic information available for *A. wentii* to facilitate the identification of terpenoid biosynthetic genes that may be involved in the production of bioactive molecules. Long-read genome sequencing of *Aspergillus wentii* CBS 141173 was conducted using Oxford Nanopore Technologies (ONT) MinION MK1C. In addition, paired-end stranded RNA-seq data from two time points, 7 days and 30 days, was used for functional annotation of the assembled genome. Overall, we assembled a genome of approximately 31.2 Mb and identified 66 biosynthetic gene clusters from the annotated genome. Metabolic extracts of *A. wentii* were analysed and the production of the bioactive terpenoid asperolide A was confirmed. We further mined the assembled and annotated genome for BGCs involved in terpenoid pathways using a combination of antiSMASH and local BlastP and identified 16 terpene synthases. Phylogenetic analysis was conducted and allowed us to establish relationships with other characterised terpene synthases. We identified two terpene clusters potentially involved in pimarane-like diterpenoid biosynthesis. Finally, the analysis of the 16 terpene synthases in our 7-day and 30-day transcriptomic data suggested that only four of them were constitutively expressed under laboratory conditions. These results provide a scaffold for the future exploration of terpenoid biosynthetic pathways for bioactive molecules in *A. wentii*. The terpenoid clusters identified in this study are candidates for heterologous gene expression and/or gene disruption experiments. The description and availability of the long-read genome assembly of *A. wentii* CBS 141173 further provides the basis for downstream genome analysis and biotechnological exploitation of this species.

## BACKGROUND

*Aspergillus* is one of the best studied genera of filamentous fungi, largely because of the medical and industrial relevance of some of its species. Their ubiquitous nature enables them to perform roles as saprophytes, parasites and endophytic symbionts with other entities in the environment, arguably inspiring the diverse secondary metabolic pathways exhibited by these organisms that range from toxins to antimicrobial molecules [1]. These secondary metabolites can be classified into various groups such as polyketides, terpenoids, peptides, alkaloids and meroterpenoids, which have raised pharmaceutical and agrochemical interests for human use. A noteworthy example is the cholesterol-lowering agent lovastatin produced by *Aspergillus terreus* [2]. For context, in the past 40 years, natural products and their derivatives from living organisms have accounted for 32% of the total small molecule drugs approved by the U.S. Food and Drug Administration with major impact as anticancer and antibacterial preparations [3].

Within the Aspergillus genus, *Aspergillus wentii* is known to produce a plethora of bioactive molecules, including 97 reported compounds that include terpenoids, anthraquinones and xanthones [4]. To understand and exploit the biosynthesis of bioactive molecules, genome sequencing is essential.

The genome sequence of *Aspergillus wentii* CBS 141173 was first reported in 2017, as part of a study conducted by a global consortium of mycologists to provide better genomic coverage for the genus *Aspergillus* [5]. However, only short-read sequencing was employed in this case.

In the present study, we explored *Aspergillus wentii* CBS 141173, a commonly found fungus in terrestrial, marine and endophytic environments and a talented producer of small molecules. This strain is a promising reservoir of a specific group of natural products with valuable biological potentials for application in medicine known as norditepenoid dilactones, which include asperolide A, wentilactone A and B (Figure 1) [6,7]. They belong to a family of diterpenoids that includes tetranorlabdane, podolactone, aromatic norditerpenes, and isopimarane. Most importantly, wentilactone A has been reported to exert a significantly selective toxicity by its inhibitory effect on lung carcinoma cell lines [8]. A recent review by Ainousah *et al*. extensively discussed *A. wentii*, highlighting the terpenoids produced by this organism [4]. This group of compounds displayed various biological properties including *in vitro* and *in vivo* antifungal, cytotoxic, herbicidal, and plant-growth-regulating activity.

**Figure 1:**
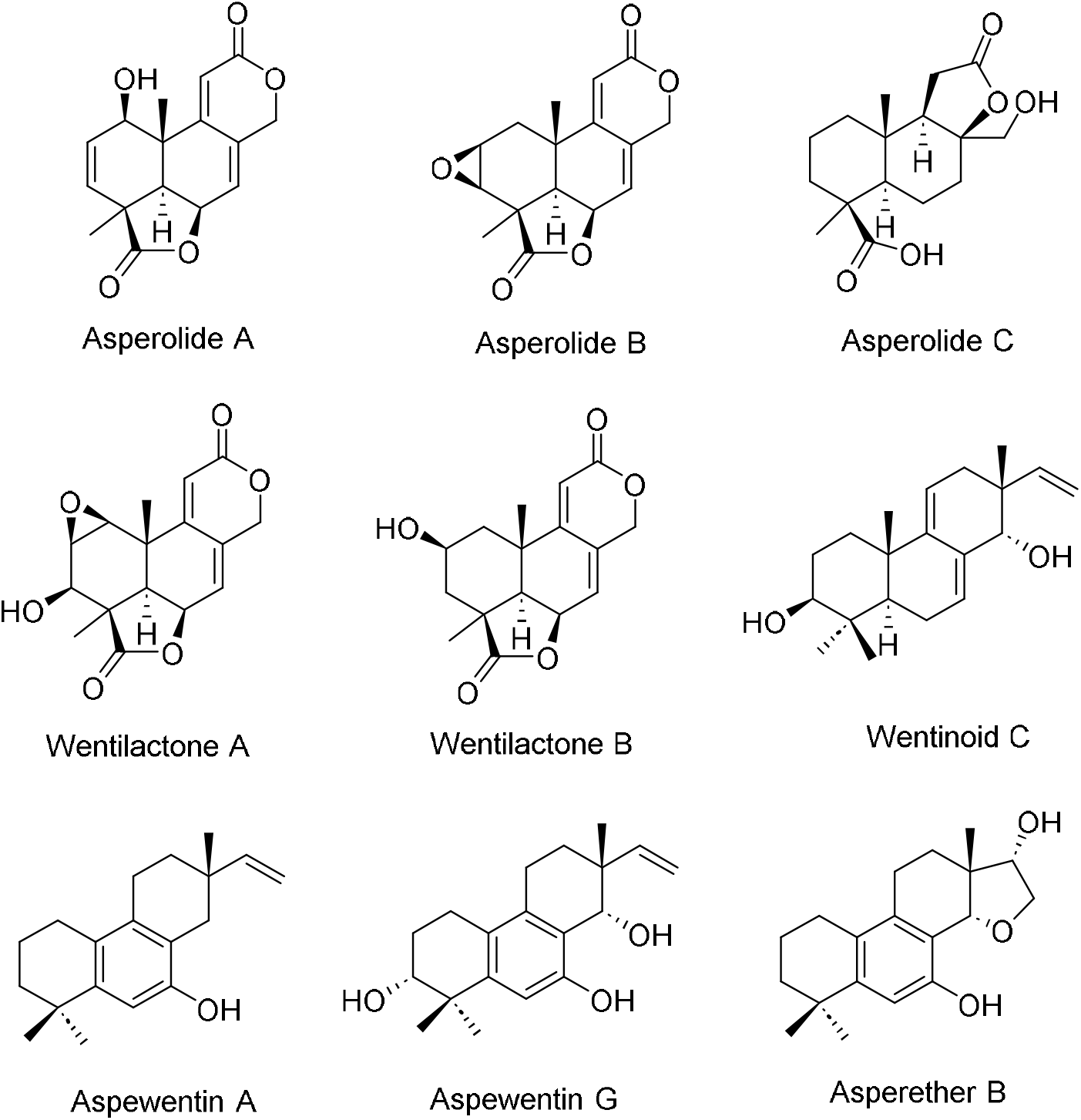
Examples of natural products reported from *Aspergillus wentii*.

Fungal terpenoid natural products are derived from the common five-carbon isoprene units isopentenyl diphosphate (IPP) and dimethylallyl diphosphate (DMAPP), which are synthesized from the mevalonate (MVA) pathway [9]. These isoprene units can be joined together to form various terpenoid scaffolds by the catalytic activity of terpene synthases; geranyl pyrophosphate (GPP) synthase, farnesyl pyrophosphate (FFP) synthase and geranyl geranyl pyrophosphate (GGPP) synthase leading to monoterpenes (C10), sesquiterpenes (C15) and diterpenes (C20), respectively [10]. It has been proposed that the family of norditerpenoids could derive from the removal of four methylene groups from a C20 diterpenoid, therefore producing a C16 molecule [11].

Clustering of genes that collaborate towards biosynthesis of secondary metabolites is a genomic phenomenon that has been widely recognised [12]. They are often described as biosynthetic gene clusters (BGCs) and have greatly aided researchers in characterising and even directing biosynthesis of the corresponding metabolite [13]. BGCs can be searched for within assembled genomes using a range of bioinformatics tools such as antiSMASH [14], cblaster [15] and local Blast searches.

In the present work, we aimed to complement the existing short-read genome assembly of *A. wentii* CBS 141173 published by de Vries *et al*. [5], with a nanopore long-read assembly and functional annotation framework of paired-end stranded RNA-seq data. The newly annotated genome and transcriptomic data will aid the elucidation of the terpene synthase biosynthetic gene clusters that could putatively be involved in the production of norditerpenoid lactone secondary metabolites and in the characterisation of other cryptic BGCs present within the genome.

## MATERIALS AND METHODS

### Fungal strain

*Aspergillus wentii* CBS 141173 was obtained from CBS-KNAW, Fungal Biodiversity Centre, Westerdijk, Netherlands. The strain was grown and maintained on Potato Dextrose Agar (PDA, 20 g/L glucose, 4 g/L potato extract, 20 g/L agar) plates and Potato Dextrose Broth (PDB, 20 g/L glucose, 4 g/L potato extract).

### Genomic DNA extraction

*A. wentii* CBS 141173 cultures were grown on 25 mL PDB within 250 mL conical flasks, using agar plugs from PDA plates containing the fungus as the inoculum. The flasks were incubated for 72 hours, at 25°C, shaking at 200 rpm. Fungal mycelia were pelleted from the broth via centrifugation at 8,000 g for five minutes, the pellet was subsequently washed twice with distilled water to remove residual media. High Molecular Weight (HMW) genomic DNA extraction was carried out with via cryogenic grinding with liquid nitrogen, and a sterile mortar and pestle. 1 g of fresh mycelia was grinded into a fine powder. 200 mg of the powder were transferred aseptically into a microcentrifuge tube.

For genomic DNA extraction, GenElute™ Plant Genomic DNA Miniprep Kit (Sigma-Aldrich) was used, following the manufacturer’s protocol. Extracted genomic DNA was further concentrated by ethanol precipitation. Briefly, 1/10 volume of 3 M sodium acetate, pH 5.2, was added to the eluted gDNA, followed by three volumes of 100% ethanol. The samples were mixed by gentle inversion and stored at −20 °C overnight, then spun at 14,000 g for 30 minutes at 4°C. The supernatant was decanted, the DNA pellet was air dried and resuspended in nuclease free water (ThermoFisher).

To ascertain the quality and concentration of extracted genomic DNA, gel electrophoresis (1% w/v agarose gel with 1 x GelRed), Nanodrop ND 1000 (Thermo Fisher, USA), and Qubit (dsDNA Broad range assay, Invitrogen) were used.

### RNA extraction and sequencing

*A. wentii* CBS 141173 total RNA was extracted using Plant/Fungi total RNA purification kit (Norgen Biotek) following the manufacturer’s instructions, based on column purification technology. An agar plug of growing mycelium was taken from a PDA plate and used to inoculate 50 mL of PDB in a 250 mL flask. Collection of the mycelia at two timepoints was performed, after 7 days and 30 days of growth, respectively, at 25°c with shaking at 200 rpm. The mycelium was harvested by centrifugation at 7,000 x g for 3 minutes, supernatant discarded, and 1 g of mycelium was grinded into a fine powder in liquid nitrogen using sterile mortar and pestle in an RNAse decontaminated environment. Using a sterile inoculation loop, 100 mg of the fine powder were transferred into a sterile 1.5 ml microcentrifuge tube and kept on dry ice. The RNA extraction and DNAse treatment were performed following the manufacturer’s instruction. The purified RNA was visualised on a 1% w/v agarose gel with 1x GelRed to assess integrity. Quantification of the RNA and purity checks were conducted with Nanodrop ND 1000 (Thermo Fisher). The RNA was sequenced with Illumina paired-end (2×150 bp) technology on a HiSeq 2500 platform (Azenta Life Sciences).

### Genome sequencing

The genome of *A. wentii* was sequenced with Oxford Nanopore Technology (ONT)(16). The manufacturer’s protocol for the Ligation sequencing kit (SQK-LSK109) was followed with minor modifications introduced for DNA repair and end-prep stage by increasing the incubation time and temperature after washing with ethanol to 15 minutes at 37°C. To enrich DNA fragments of 3 kb or longer, the Long Fragment Buffer was used for the DNA clean-up. Noteworthy, 1.2 µg of genomic DNA was used for the library preparation, resulting in a final amount of 345 ng of DNA at the end of the adapter ligation and clean-up step. Nanopore sequencing was performed on a MinION MK1C Platform with a FLO-MIN-106 R9.4 flow-cell and run for 18 hours.

### Genome assembly

*De novo* strategies were utilised for the fungal genome assembly. Utilizing the default parameters, raw data from the MinION MK1c output .fastq files were transferred and merged into one single .fastq file and inputted into Flye version 2.8.2 using default settings and the nano-raw mode, to create a draft genome assembly. Polishing of this draft assembly was done by aligning the merged .fastq file that was inputted into Flye against the draft genome using Minimap2 version 2.11, errors could then be corrected using Racon version 1.4.20 [17]. The settings used in Racon were as follows: a value of 8 for matching bases, −6 for mismatches, −8 for gaps, and a window size of 500. Polishing using this procedure was performed three times. The third Racon polished assembly was then inputted into Medaka version 1.2.1 (https://github.com/nanoporetech/medaka) using model r941_min_high_g360 and default settings creating a consensus fasta file.

Polishing was then performed on the consensus file generated by medaka using Pilon version 1.24 [18] and its default settings. The output from Pilon was polished again, this was repeated until Pilon had been run four times. For the polished genome evaluation, Benchmarking Universal Single-Copy Orthologs (BUSCO) version 5.6.1 was used with eurotiales_odb10 database with the default genome mode settings [19].

### Genome annotation

Functional annotation was done using the Funannotate version 1.8.17 pipeline [20], utilizing a “gold standard” annotation framework of paired-end stranded RNA-seq data from two time points, 7 days and 30 days coupled with a genome assembly built from a *de novo* strategy, following the Funannotate user instructions of clean-sort-mask. The following step of Funannotate training involved the introduction of RNA sequencing data, using downstream commands Funannotate predict and Funannotate update. Results generated were then run through antiSMASH v7.1.0 [14] within the Funannotate pipeline. The final prediction was run on BUSCO [19] to provide quality check on the predicted proteins generated, once again using the eurotialales_odb10 database. All parameters not mentioned above were left on their default setting, unless stated otherwise. The annotated genome of *A. wentii* CBS 141173 was further analysed by running EggNOG-mapper v2.1.12 [21] and programs that are part of the GenomeTools 1.6.1 package [22].

### Fungal metabolite extraction

*A. wentii* CBS 141173 was cultured on PDA plates at 28°C for 10 days and then agar plus were used to inoculate 100 mL of PDB medium in 500 mL flasks, which were grown for 30 days at 25 °C, shaking at 200 rpm. The culture was then homogenised by TissueRuptor (Qiagen) to a fine consistent paste. The blended mix was acidified with concentrated hydrochloric acid to pH= 3. Thereafter, it was twice extracted with ethyl acetate with a ratio of 1:1 v/v. The organic phases resulting from the two extractions were pooled, dried with addition of an excess of MgSO_4_, filtered and evaporated *in vacuo*. The resulting residues were dissolved in high-performance liquid chromatography (HPLC)-grade methanol, spin-filtered, transferred to amber glass 11 mm vials and analysed by Ultra-high performance Liquid Chromatography-High-Resolution Mass Spectrometry (UHPLC-HRMS).

### UHPLC-HRMS Analysis

UHPLC-HRMS analyses were carried out on 5 μL of prepared extracts injected through a reverse phase column using an UHPLC-HRMS Agilent ZORBAX Eclipse Plus column C18, 4.6 × 150 mm 40 (particle size 5 μm) on a Dionex 3000RS UHPLC coupled to a Bruker Ultra High Resolution (UHR) Q-TOF MS MaXis II mass spectrometer with an electrospray source. Sodium formate (10 mM) was utilised for internal calibrations, with a gradient elution from 95 : 5 solvent A/solvent B to 0 : 100 solvent A/solvent B in 35 minutes, with solvent A being water (0.1% formic acid) and solvent B acetonitrile (0.1% formic acid).

### Identification of terpene synthases

The long-read genome assembly of *A. wentii* incorporating full length transcriptomic RNA-seq data was used in this study. This allowed the identification of putative terpene synthase (TS) gene family based on antiSMASH predictions and local BlastP analysis using as a query TS genes from NCBI GenBank (https://www.ncbi.nlm.nih.gov/genbank/) and UniProt (https://www.uniprot.org/).

### Phylogenetic analysis

Multiple sequence alignments of protein sequences of mined TSs and their homologues from other organisms were conducted using MUSCLE in MEGA11.0 using default settings. The obtained alignment was used as the input for the Maximum Likelihood tree algorithm in MEGA11.0 software to construct a phylogenetic tree [23]. Visualisation was conducted using iTOL: Interactive Tree Of Life web viewer https://itol.embl.de/.

## RESULTS

### Genomic DNA and total RNA extraction

High molecular weight (HMW) genomic DNA of *Aspergillus wentii* was sequenced using Oxford nanopore long-read whole-genome sequencing. HMW genomic DNA extraction was performed on *A. wentii* in duplicate (Supplementary Figure 1). The extracted genomic DNA was further subjected to ethanol precipitation and resuspended in 35 µL of nuclease-free water leading to an improved concentration of both samples (Supplementary Table 1). Fungal total RNA was extracted in duplicate from *A. wentii* liquid cultures at two time points, 7 days and 30 days, and sequenced using Illumina paired-end (2×150 bp) sequencing. The number of RNA-seq reads obtained for each sample is shown in Supplementary Table 2.

### Genome assembly and analysis

The nanopore sequencing generated a significant quantity of long-read data with a high degree of coverage of over 100x. The large coverage of the long-read nanopore sequencing allowed for a robust first draft genome to be created, with only few fragments. To improve the annotation framework, we included paired-end RNA-seq data from two time points, which is known to lead to a high-quality annotation facilitated by Funannotate 1.8.17 [20].

The final draft genome size for the organism was 31.2 Mb. This is consistent with the size of other Aspergilli genome assemblies which tend to be 31–40 Mb in size [5,24]. The full genome statistics (Table 1), were generated with genometools [22]. Briefly, the genome comprised 8 contigs, 7 of which may be putatively assembled at the chromosome level and 1 being putatively mitochondrial

**Table 1:** Genome statistics and predicted features of the nanopore assembled genome of *Aspergillus wentii* CBS 141173.

| Parameters | *Aspergillus wentii* CBS 141173 |
| --- | --- |
| number of contigs | 8 |
| total contigs length | 31252061 |
| as % of genome | 156.26 % |
| mean contig size | 3906507.62 |
| contig size first quartile | 2796016 |
| median contig size | 4343183 |
| contig size third quartile | 5069049 |
| longest contig | 7905159 |
| shortest contig | 27878 |
| contigs > 500 nt | 8 (100.00 %) |
| contigs > 1K nt | 8 (100.00 %) |
| contigs > 10K nt | 8 (100.00 %) |
| contigs > 100K nt | 7 (87.50 %) |
| contigs > 1M nt | 7 (87.50 %) |
| N50 | 4417627 |
| L50 | 3 |
| N80 | 4127719 |
| L80 | 5 |
| NG50 | 5069049 |
| LG50 | 2 |
| NG80 | 4417627 |
| LG80 | 3 |

DNA. The genome has a 47.2% GC content, N50 length of 4,417,627 bp, with the longest contig being 7,905,159 bp and the average gene length being 1,365 bp. A total of 12,925 genes and 187 tRNAs were predicted from the genome.

While the gene function of the predicted genes in genomes was assigned using Funannotate, the predicted proteome was run again through Eggnog-mapper v2.1.4-2 [21]. This yielded a prediction of 10,592 proteins. Supplementary Figure 2 shows how through eggnog-mapper we were able to assign gene ontology terms to the genome including matches to multiple KEGG databases, and BRITE hierarchies. Additionally, predicted proteins were matched to the Carbohydrate Active enZymes (CAZymes) CAZy database and a total number of 1,615 genes were predicted to be CAZy. Noteworthy, >85% of genes in the genome were matched to proteins in the pfam database. All data generated using eggnog mapper are available in the Supplementary Dataset 1.

From the predicted proteins from eggnog-mapper, 85.7% Clusters of Orthologous Groups (COG) categories were assigned. The distribution of the predicted proteins across the COG categories is shown in Supplementary Figure 3. The most frequently mapped category was “Function Unknown”. Of those that could be placed into a category of known function, the five most common in order of decreasing predicted protein count within the categories were “Intracellular trafficking, secretion, and vesicular transport”, “Amino acid transport and metabolism”, “Secondary metabolite biosynthesis, transport and catabolism”, “Posttranslational modification, protein turnover, chaperones”, and “Carbohydrate transport and metabolism”. The presence of over 575 proteins associated with secondary metabolites in the genome is promising and these deserve further examinations, particularly in the context of the high value terpenoids such as wentilactones, and asperolides known to be produced by *Aspergillus wentii*. It is also interesting that no mobilome elements associated with transposons and prophages were detected, and less than 100 predicted proteins were identified in the genome that were associated with any of the following categories: “Extracellular structures”, “Cell motility”, “Nuclear structure”, and “Defence mechanisms”.

Finally, BUSCO was used to assess the completeness of the genome assembly and annotation using the Eurotiales odb10, and suggested a complete annotation set with a 94.8% genome BUSCO score and a 95.3% protein BUSCO score, respectively (Supplementary Table 3).

### Biosynthetic gene clusters analysis

The whole genome sequence of *Aspergillus wentii* produced in this study, as well as the one produced previously through short-read sequencing [5] were mined for putative biosynthetic gene clusters (BGCs) using antiSMASH [14]. A total number of 66 BGCs were identified from the sequenced genome (Supplementary Dataset 2). This is an improvement compared to 50 BGCs identified from the genome previously deposited to NCBI and JGI (Table 2). From the present study, 19 non-ribosomal peptide synthetases (NRPS) BGCs were found to be the most abundant in the genome, followed by polyketide synthases (PKS) and terpene BGCs.

**Table 2:**
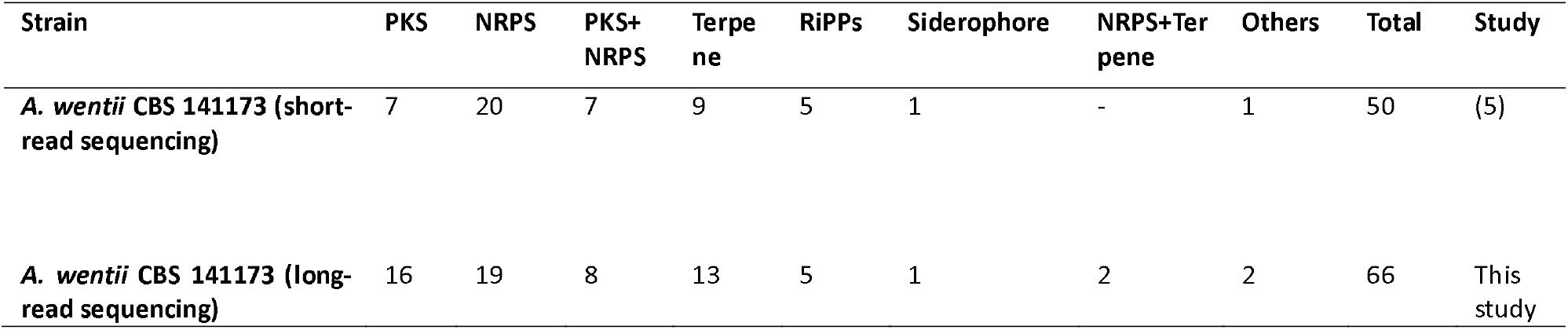
Comparative antiSMASH analysis of predicted BGCs from the two *Aspergillus wentii* CBS 141173 genome assemblies.

### Comparative genomics

To further analyse the biosynthetic capabilities of *Aspergillus* genomes, a BGC comparative analysis was conducted using the BGC identification defined by antiSMASH (Figure 2). Across the set genomes used for this analysis, it is noteworthy that *Aspergillus parasiticus* contained the highest number of BGCs - 77. Interestingly, the highest number of putative terpene BGCs - 13 - was found in the long-read assembly of *A. wentii*, followed by *A. felis* and *A. nidulans* with 11 and 10, respectively. NRPS and PKS BGCs were altogether the most dominant class across the Aspergilli genomes. For instance, 26 NRPS clusters were identified from both *A. flavus* and *A. parasiticus*. Our long-read assembly of *A. wentii* recorded a high number of polyketide BGCs - 16 - while the highest was 17 from the *A. parasiticus* genome. There is a vast resource of fungal genomes to always inspire BGCs comparative studies, notwithstanding, our BGC analysis shows that the long-read assembly of *A. wentii* CBS41173 is consistent with other *Aspergillus* genomes in its capacity to produce diverse natural products.

**Figure 2:**
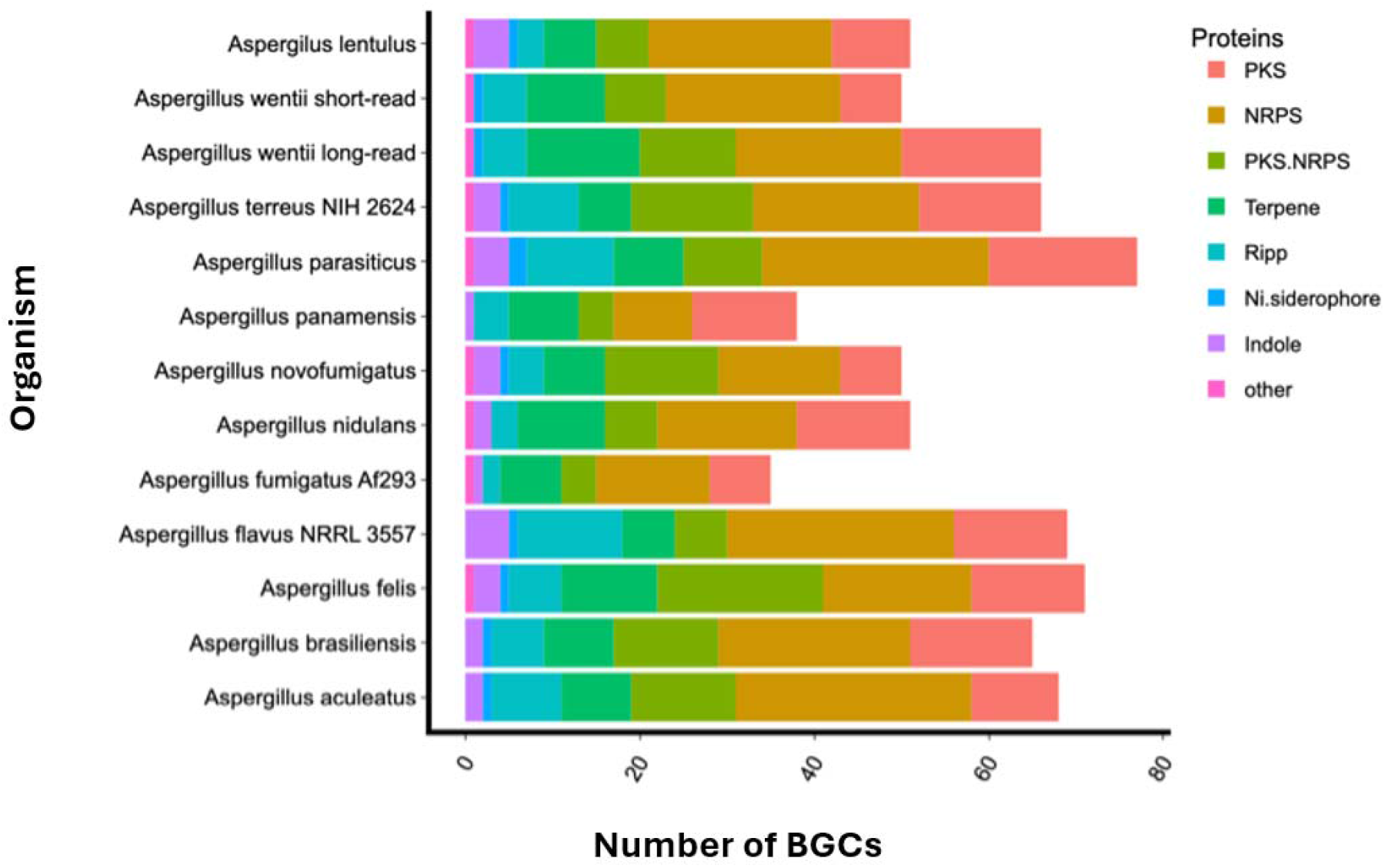
Comparison of the BGCs found in selected *Aspergillus* genomes. The number of BGCs from the sequenced *A. wentii* fungal strain was compared to the number of BGCs from other members of the *Aspergillus* genus as predicted by antiSMASH. This revealed that the fungal strain characterised in this study has the potential to produce terpenoids, polyketides and NRPs in line with other members of the genus.

### Production of asperolide A

UHPLC-HRMS was used to analyse metabolite extracts from *A. wentii* CBS 141173. Asperolide A was detected from the metabolic extracts on the basis of the calculated m/z 289.1076 [M+H]^+^ for asperolide A. Extracted ion chromatogram of the HRMS data showed a dominant ion peak at m/z 289.1072 (Figure 3) and the molecular formula was predicted as C_16_H_16_O_5_, confirming the production of the norditerpenoid compound. Speculatively, asperolide A might be the main norditerpenoid produced by *A. wentii* CBS 141173 at the 30-day time point analysed under laboratory conditions.

**Figure 3:**
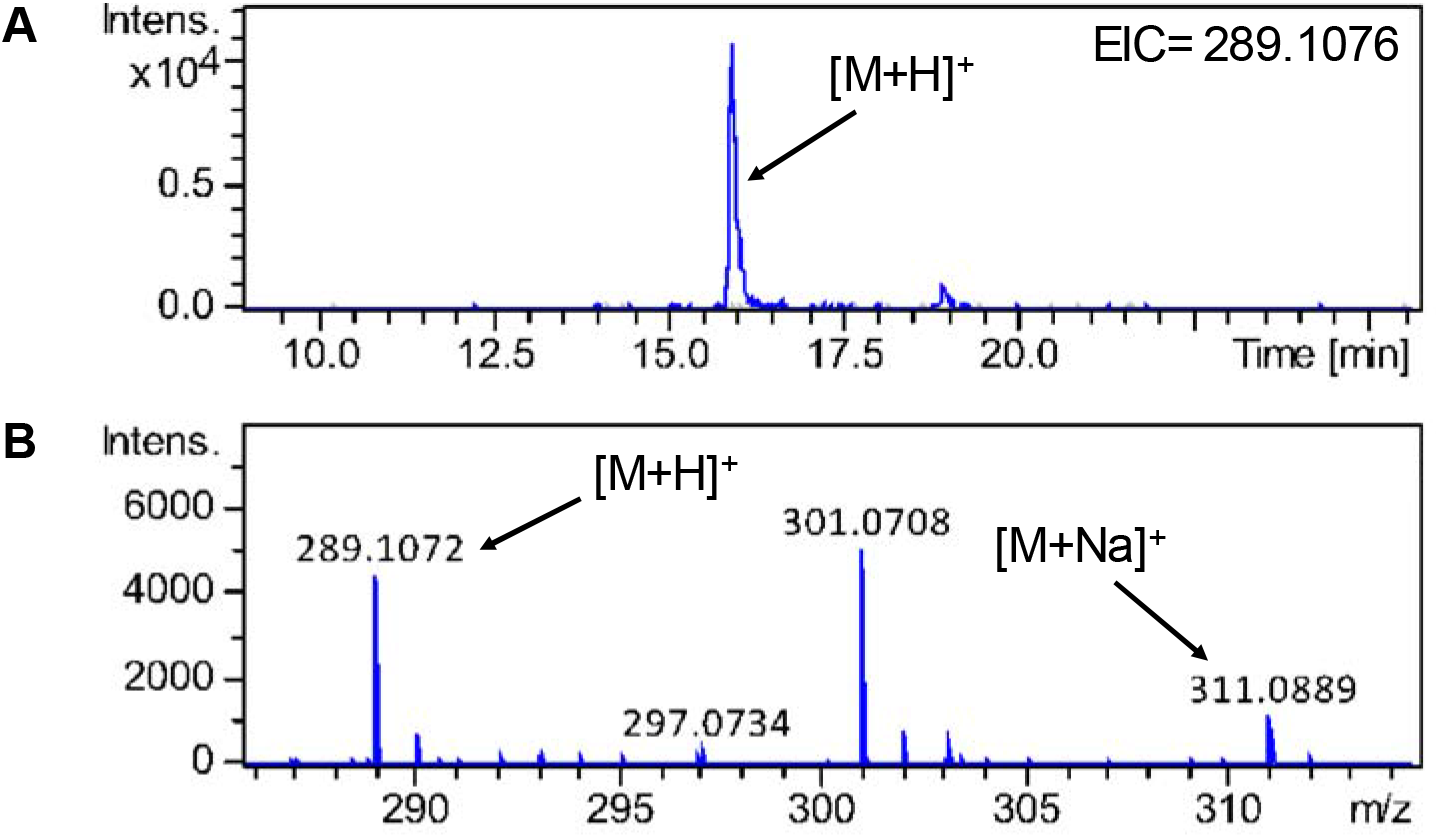
UHPLC-HRMS detection of asperolide A. **A**) Extracted ion chromatogram in positive mode for m/z = 289.1076 is shown, highlighting accumulation of asperolide A in A. wentii (trace in blue) and not in the culture medium control (trace in grey). **B**) Mass spectrum of asperolide A (M= C_16_ H_16_ O_5_); m/z expected for [M + H]^+^= 289.1076, for [M + Na]^+^= 311.0895.

### Terpene synthases from Aspergillus wentii CBS 141173

To identify putative terpene synthases that could be involved in the production of terpenoids in A. wentii, we utilised two strategies, antiSMASH and local BlastP analysis. Noteworthy, from the 13 putative terpenoid clusters identified by antiSMASH, we shortlisted 10 (AwTS1 – AwTS10), based on their protein blast (BlastP) homology with other known TSs from the UniProt database (Table 3). Using a manual BlastP analysis of the Aspergillus wentii BGCs on antiSMASH, we also recruited additional terpene synthases (AwTS11 – AwTS14) found within other BGCs such as polyketide and NRPS clusters. Finally, a local BlastP search using geranylgeranyl pyrophosphate synthase as a query allowed us to identify pimarane/pimaradiene-like synthases (AwTS15, AwTS16) from the genome assembly.

**Table 3:** BlastP analysis of terpene synthases from *Aspergillus wentii* CBS 141173 genome.

| Name | GeneID | Homologue (% identity/% similarity), Organism | Homologue accession | Method of Identification |
| --- | --- | --- | --- | --- |
| **AwTS1** | AWENTII_000319 | Squalene hopane cyclase *afumA* (50/64) *Aspergillus fumigatus* A1163 | B0Y565.1 | antiSMASH |
| **AwTS2** | AWENTII_001688 | bicyclogermacrene synthase (31/45) *Penicillium expansum* | A0A0A2JP58.1 | antiSMASH |
| **AwTS3** | AWENTII_002622 | Lanosterol synthase *erg7A* (89/92) *Aspergillus fumigatus* Af293 | Q4WES9.1 | antiSMASH |
| **AwTS4** | AWENTII_003014 | Terpene cyclase (24/42) *Fusarium fujikuroi* IMI 58289 | S0EGZ9.1 | antiSMASH |
| **AwTS5** | AWENTII_006486 | Lanosterol synthase *erg7A* (41/59) *Aspergillus fumigatus* Af293 | Q4WES9.1 | antiSMASH |
| **AwTS6** | AWENTII_006746 | Bifunctional lycopene cyclase (59/66) *Nannizzia gypsea* CBS 118893 | E4UPP6.1 | antiSMASH |
| **AwTS7** | AWENTII_009676 | (E)-beta farnesene synthase MBR_03882 (32/52) *Metarhizium brunneum* ARSEF 3297 | A0A0B4G504.1 | antiSMASH |
| **AwTS8** | AWENTII_009792 | Squalene hopane cyclase *afumA* (44/58) *Aspergillus fumigatus* A1163 | B0Y565.1 | antiSMASH |
| **AwTS9** | AWENTII_010168 | Geranylgeranyl pyrophosphate synthase (66/82) *Neurospora crassa* OR74A | P24322.2 | antiSMASH |
| **AwTS10** | AWENTII_012038 | Farnesyl-diphosphate farnesyltransferase *erg9* (79/88) *Aspergillus fumigatus* Af293 | Q4WAG4.1 | antiSMASH |
| **AwTS11** | AWENTII_000061 | Terpene cyclase *dpasB* (38/58) *Apiospora sacchari* | P9WEX1.1 | antiSMASH |
| **AwTS12** | AWENTII_005314 | Fusicoccadiene synthase (29/47) *Diaporthe amygdali* | A2PZA5.1 | antiSMASH |
| **AwTS13** | AWENTII_005350 | Terpene cyclase *flvE* (34/50) *Aspergillus flavus* NRRL3357 | B8NHE0.1 | antiSMASH |
| **AwTS14** | AWENTII_006388 | (E)-beta farnesene synthase MBR_03882 (35/54) *Metarhizium brunneum* ARSEF 3297 | A0A0B4G504.1 | antiSMASH |
| **AwTS15** | AWENTII_006883 | Pimaradiene synthase *pbca* (61/75) *Aspergillus nidulans* FGSC A4 | A0A1U8QHE3.1 | Local BlastP |
| **AwTS16** | AWENTII_011959 | Pimaradiene synthase *pbca* (43/60) *Aspergillus nidulans* FGSC A4 | A0A1U8QHE3.1 | Local BlastP |

### Terpene biosynthetic gene clusters

To understand the genetic context of the terpene synthases identified, we analysed the flanking regions of each of the terpene synthase genes. Full descriptions of the gene clusters can be found in Supplementary Dataset 3. Worth highlighting is that the AwTS15 and AwTS16 terpene clusters are predicted to encode a pimarane/pimaradiene diterpenoid scaffold. For the AwTS15 BGC (Figure 4A), BlastP analysis shows the presence of primary metabolic enzymes such as the HMG-CoA involved in the mevalonate pathway that may be involved in the biosynthesis of geranylgeranyl pyrophosphate to make a diterpenoid compound. The additional presence of multifunctional biosynthetic enzymes such as oxidoreductase and a cytochrome P450 in the cluster could potentially be involved in the maturation of the diterpenoid scaffold. Also, for the AwTS16 BGC (Figure 4B), the presence of three cytochrome P450s and other tailoring enzymes makes it a candidate cluster for the production of pimarane-like diterpenoids biosynthesis in *A. wentii*. Other notable terpenoid clusters found in the genome include the AwTS1 triterpenoid cluster (Figure 4C) and AwTS13 hybrid terpene cluster with an NRPS biosynthetic origin (Figure 4D). Experimental characterisation will be vital to elucidate the resulting biosynthetic products.

**Figure 4:**
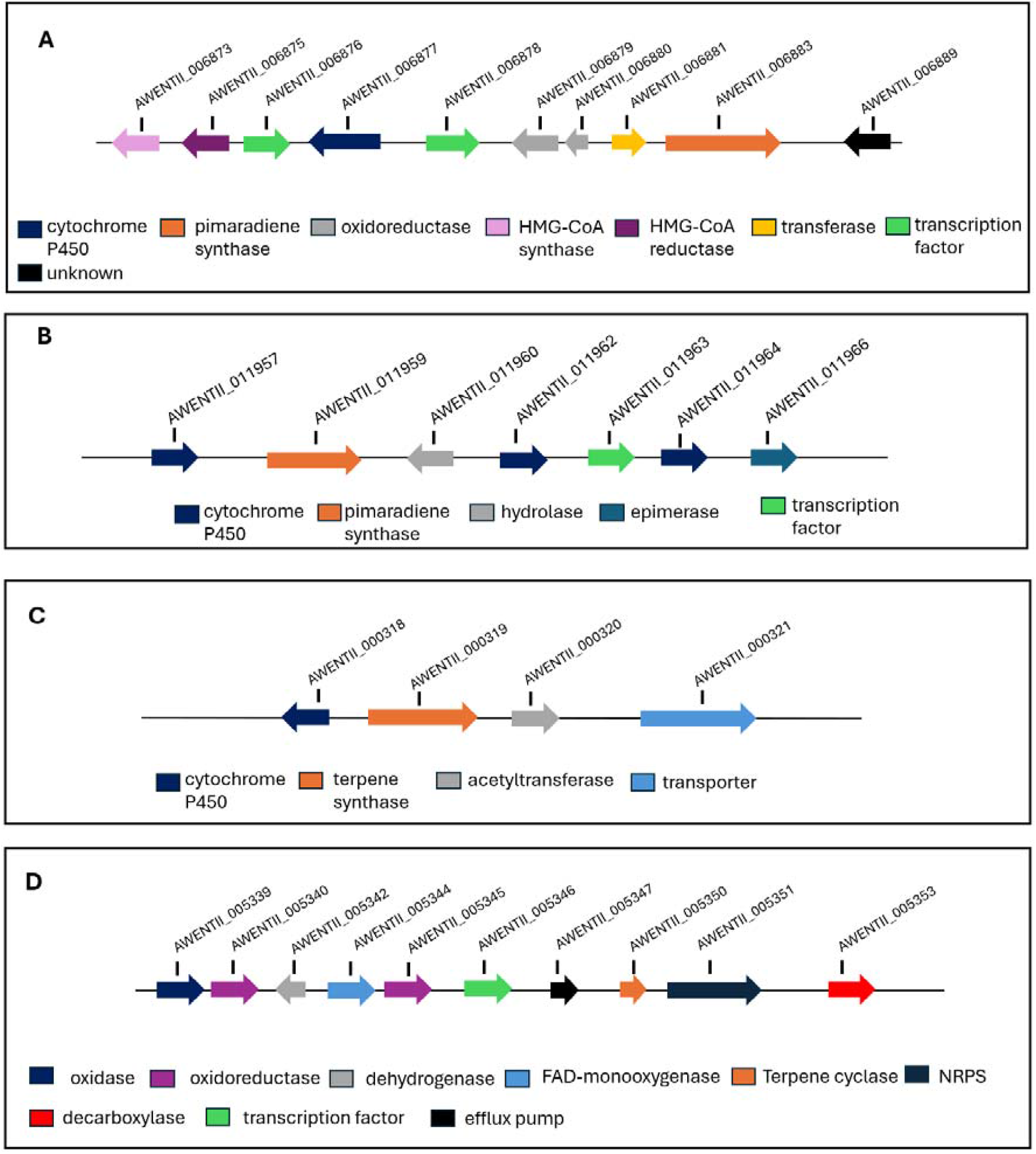
Diversity of the terpenoid biosynthetic gene clusters from *A. wentii* CBS 141173. **A**) AwTS15 diterpenoid gene cluster. **B**) AwTS16 diterpenoid gene cluster. **C**) AwTS1 triterpenoid gene cluster, **D**) AwTS13 hybrid terpene-NRPS biosynthetic gene cluster.

### Terpene Phylogeny

To further investigate the role of the terpenoid genes from *Aspergillus wentii* in making terpenoid compounds, a phylogenetic assessment was also conducted using 55 TS proteins, comprising the 16 identified AwTSs from *Aspergillus wentii* (Table 3), and 39 characterised TSs from other fungal species (Supplementary Dataset 4). These terpene synthases were grouped into subfamilies (TFa-TFj) based on their putative role in terpenoids biosynthesis such as diterpenoid pyrones, tricyclic diterpenoids, and mixed terpenoids (Figure 5).

**Figure 5:**
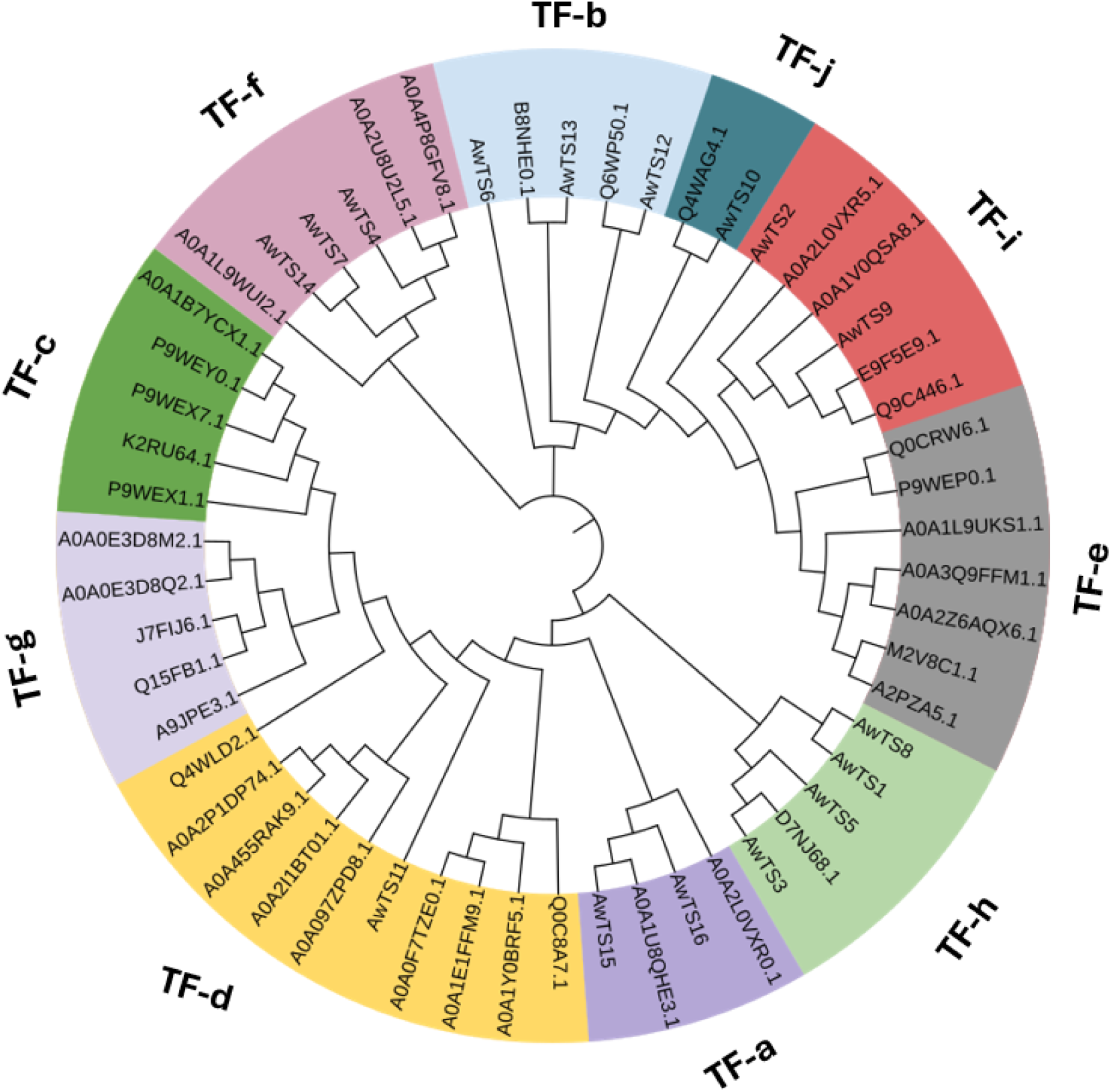
Phylogenetic analysis of *Aspergillus wentii* CBS 141173 terpene synthases (AwTS), based on their predicted protein sequences. The Maximum Likelihood algorithm tree was generated from an alignment of 55 TS proteins, comprising 16 AwTS and another 39 characterised TSs from different fungal species. Sections are identified by colours. The names of the sections are indicated as **TF-a**: Tricyclic diterpenoids, **TF-b**: Mixed diterpenoids, **TF-c**: Diterpenoid pyrones, **TF-d**: Mixed sesquiterpenoids, **TF-e**: Sesterterpenoids, **TF-f**: α-humulene based terpenoids, **TF-g**: Indole diterpenoids, **TF-h**: Squalenes, **TF-i:** Diterpenoids, **TF-j:** Lanosterol transferases.

From the phylogeny analysis, the *A. wentii* terpene synthases appeared to be very diverse and share similarities with some bifunctional cyclases (TF-a, and TF-f) such as those that could encode for terpenoid scaffolds such as monocyclic humulene-like compounds and tricyclic diterpenoids, respectively. Additionally, we observed relationships between AwTS6, -11, -12 and -13 and terpene synthases involved in mixed biosynthetic pathways (TF-b and TF-d) (Figure 5). AwTS2 and AwTS9 fell into the TF-i subfamily, revealing their relationship with other diterpenoid synthases. The AwTSs in the TF-h section are putative triterpenoid synthases and distinctly AwTS10 belong to the lanosterol transferase category. Interestingly none of the AwTS belonged to the subfamilies of diterpenoid pyrone, sesterterpenoid and indole diterpenoids.

### Expression of AwTSs under laboratory conditions

We further analysed the count of the RNA transcripts of each of the terpene genes by comparing them with the beta-tubulin and GAPDH (housekeeping) genes of *A. wentii* to gain insights into the level of their gene expression (Table 5). Interestingly, from the 16 TSs recruited for our analysis across the two different timepoints of 7 days and 30 days, only four genes (AwTS3, -5, -9, and -10) were expressed at considerable levels, while others were suppressed or not expressed under laboratory conditions (Table 5). AwTS3 and AwTS5 are predicted to be lanosterol synthases, whereas AwTS9 and AwTS10 are predicted to be a geranylgeranyl pyrophosphate synthase and a farnesyl-diphosphate farnesyltransferase, respectively (Table 3). Moreover, AwTS9 was differentially expressed across the two timepoints, with a substantial increase in expression at 30 days. Generally, most terpenoid genes exhibited the same suppression characteristic across the two timepoints suggesting that some genes responded in similar ways to standard laboratory conditions.

**Table 5:**
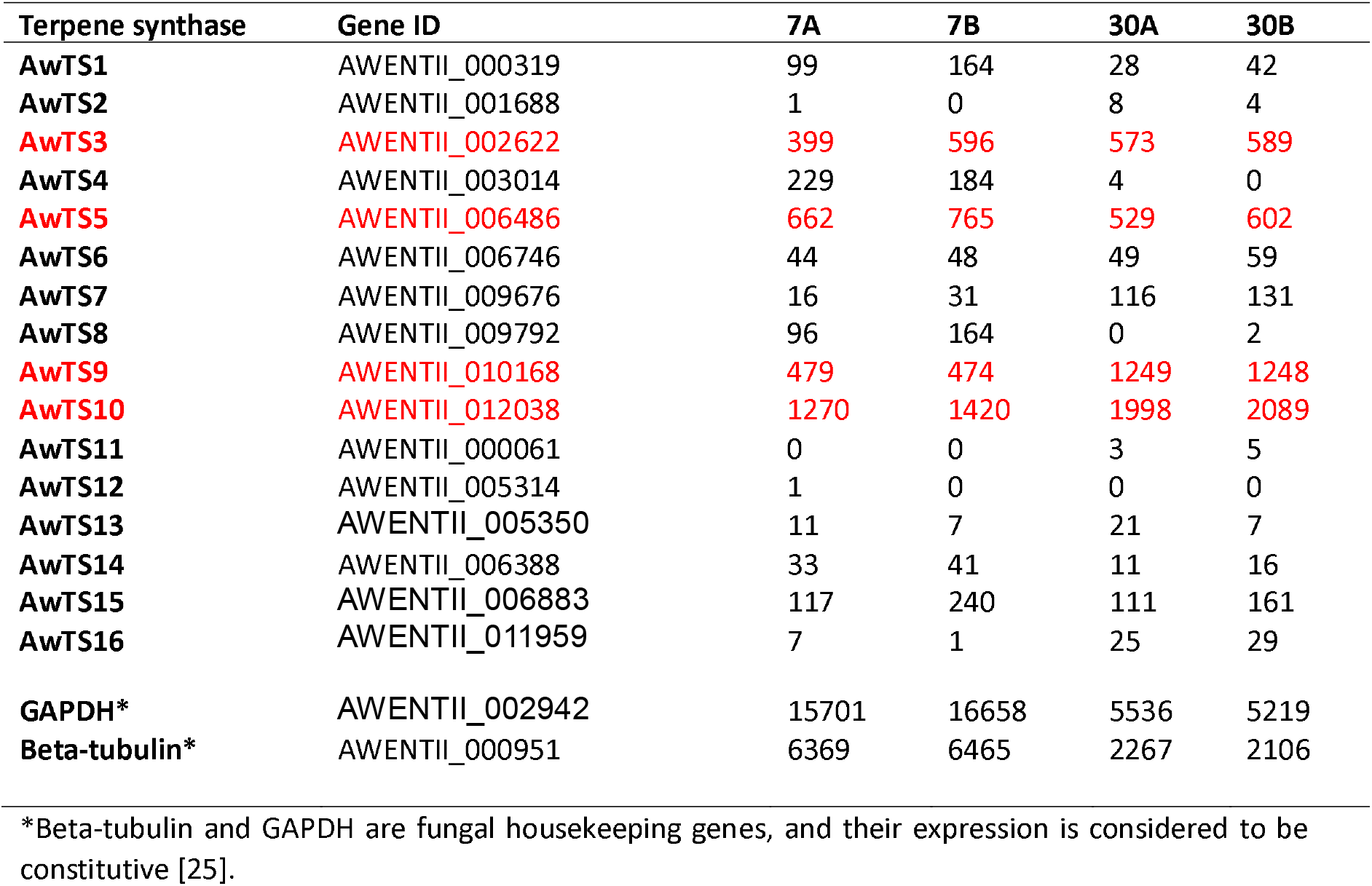
Transcriptomic hitcounts of terpenoid genes in *Aspergillus wentii* CBS 141173 at two timepoints (7 and 30 days) performed in duplicate (A and B). Genes highlighted in red (AwTS3, -5, -9, and -10) show the highest expression across the samples analysed.

## DISCUSSION

In this study, we provided the long-read genome assembly of *A. wentii* CBS 141173 with RNA-seq functional annotation, which has improved the quality of genomic resources available to carry out biosynthetic studies on this fungal strain. We successfully sequenced the whole genome of *A. wentii* to about 31.2 Mb, which is consistent with the genome sequence and assembly that was initially published [5]. The improved annotated genome assembly allowed for a significant improvement in the identification of 66 BGCs, including 16 terpene synthases, from the genome that could direct the biosynthesis of novel and known bioactive compounds. Our study shows how Nanopore sequencing can continue to spur long-read assembly of fungal genomes for targeted biosynthetic studies and is becoming increasingly useful even for both genomics and metagenomic research [26,27]. Various high-quality fungal genomes have already been sequenced on a MinION [28–30] thereby contributing to the availability of fungal genomes across curated databases such as NCBI and JGI.

Nanopore sequencing presents challenges with accuracy when compared to established short-read platforms such as Illumina [31]. The proposed standard for most whole-genome sequencing projects to achieve high quality assemblies would be a hybrid assembly by incorporating both nanopore long-read and short-read sequencing [32] as seen in Tamizi *et al*. [32], as it combines the advantages provided by the two sequencing platforms. Equally, we believe that a cost-effective stand-alone long read sequencing strategy can become a viable alternative. Providing necessary software upgrades for genome polishing tools and base calling algorithms would improve assembly confidence for downstream genomic studies [33]. Additionally, incorporating paired-end RNA-seq data to guide functional annotation of long-reads assembly can boost the integrity of the whole-genome assembly [34].

From the metabolite analysis conducted in our study, the bioactive norditerpenoid asperolide A made by *A. wentii* CBS 141173 was detected. The compound has been reported to show cytotoxicity against several human tumour cell lines in preliminary screening experiments and has been reported to inhibit lung carcinoma cell proliferation [35], bone metastatic breast cancer [36], and hepatocellular carcinoma cell proliferation by targeting MAPK and TP53 [6,7,35,37]. The structural similarities between asperolide A and other high-value anticancer norditerpenoids such as wentilactones A and B [38,39] confirms its potential as a scaffold for drug development.

To uncover the biosynthesis of such high-value molecules, the biosynthetic genes need to be identified. AntiSMASH is a powerful tool that has enabled the identification of BGCs encoding high value compounds. A classic example is strobilurin – the natural product that inspired the development of the β-methoxyacrylate class of agricultural fungicides that are widely used in crop protection [40,41]. AntiSMASH enabled the identification of the strobilurin polyketide BGC from the *Strobilurus tenacellus* genome and facilitated the characterisation of its biosynthetic pathway [40]. Other traditional approaches can also be used to identify novel BGCs. For instance, in the identification of the pleuromutilin terpenoid BGC from *Clitopilus passeckerianus*, degenerate PCR was used to identify putative GGPPS and then followed by analysis of the flanking regions to reveal additional biosynthetic genes included in the pleuromutilin BGC [42,43].

With advances in sequencing technology and continuous improvements with the bioinformatic tools, such as antiSMASH [14], and cblaster [15] finding novel terpenoid BGCs has become easier. From our study, a complementary gene mining approach using antiSMASH and local BlastP searches enabled the identification of BGCs that might not be picked up by antiSMASH alone due to the lack of canonical synthase genes in some BGCs [44]. With this approach we were able to recruit 16 terpene synthases for downstream analysis.

We further analysed the flanking regions of the terpene synthases to reveal their potential to encode diterpenoid compounds. From our analysis, we highlighted two diterpenoid clusters from AwTS15 and AwTS16 that could be involved in the biosynthesis of pimarane diterpenoids such as pimaradiene [45], and acanthoic acid (46). Noteworthy, the pimarane carbon skeleton is a signature scaffold for notable diterpenoids isolated from *A. wentii* such as aspewentins and wentinoids [4]. The presence of tailoring enzymes such as cytochrome P450 and oxidoreductase enzymes in the highlighted cluster adds credibility to the potential of the cluster to direct the biosynthesis of a bioactive diterpenoid compound.

From the phylogeny analysis, *A. wentii* terpene synthases appeared to be very diverse and could be involved in directing monocyclic and tricyclic terpenoid based scaffolds. For instance, humulene based scaffolds are monocyclic sesquiterpenoids recognised for their cytotoxic and anti-inflammatory effects [47–49]. Although a common feature of some plant terpenoids [49], humulene synthases have been characterised from fungi such as *EupE*, which is involved in the biosynthesis of Eupenifeldin [50]. Our phylogenetic study further reiterates the possibility of some AwTS to encode tricyclic diterpenoids due to their relationship with pimaradiene synthases [51,52] and its resulting compounds are often known for their cytotoxic bioactivity [46,52]. Terpene synthases are very versatile, as they can be involved in making either stand-alone terpenoids or hybrid biosynthetic products, such as alkaloidal terpenoids from a mixed terpene and nonribosomal peptide biosynthetic pathway [53]. Since none of the terpene synthases from *A. wentii* have been experimentally characterised, it will be informative to understand the function of these genes *via* either heterologous expression or gene deletions.

Transcriptomic analysis showed that most terpenoid synthases of *A. wentii* are not expressed under standard laboratory conditions. The use of beta-tubulin and GAPDH as reference genes to analyse the transcript level of the terpenoid synthase genes shows low levels of expression across the 7- and 30-days timepoints. This is not unusual, as various strategies have been used to improve expression of BGCs in fungi. For instance, the manipulation of transcription regulators such as *mcrA* impacts fungal secondary metabolism especially in *A. wentii* to upregulate and trigger the expression of cryptic BGCs [54,55]. Additionally, the culture medium condition has a role in determining fungal metabolism and growth [56]. The availability and concentration of certain nutrients such as carbon source, minerals and pH can influence gene expressions and ultimately impact the production of secondary metabolites [56]. For example, the production of ascofuranone and ascochlorin, meroterpenoid compounds from *Acremonium egyptiacum*, was shown to be dependent on the culture medium [57]. Since *Aspergillus wentii* is associated with marine habitats [6], extracting RNA and metabolites from mycelia grown in culture medium supplemented with seawater could provide more insight into the production of some of its cryptic metabolites and other norditerpenoid dilactones which have been detected in *A. wentii*.

## CONCLUSIONS

Our study provides a platform for elucidating the biosynthesis of norditerpenoids and other specialised metabolites made by *Aspergillus wentii* CBS 141173. Under standard laboratory conditions, RNA-seq data from two time points of 7 days and 30 days enabled us to provide a robust annotation for our nanopore long-read assembly. Based on the secondary metabolite analysis of *Aspergillus wentii* CBS 141173, we were able to confirm the production of asperolide A. We further analysed 16 terpene synthases identified from our genome assembly that could be involved in the biosynthesis of various terpenoid scaffolds in *Aspergillus wentii*, and highlighted two clusters that could make diterpenoids such as pimarane-like compounds. Genetic engineering strategies such as heterologous expression and gene disruption experiments, including through CRISPR/Cas9, can be used to link the compounds to their respective biosynthetic genes and enable us to elucidate the terpenoid biosynthetic pathways for bioactive molecules such as asperolide A made by *A. wentii*.

## Ethics approval and consent to participate

Not applicable.

## Consent for publication

Not applicable.

## Availability of data and materials

The assembled annotated genome was deposited at the NCBI database under the BioProject: PRJNA1133277, accession number: CP165651-CP165658 (Supplementary Table 4). RNA-seq raw data is available upon request.

## Competing interests

None.

## Funding

F.A. was supported by a UKRI Future Leaders Fellowship [MR/V022334/1]. Authors’ contributions F.A., C.C. and R.O. were responsible for project conceptualisation. R.O. and F.A. were responsible for experimental design and analysis. R.O. analysed and interpreted the data. R.O. wrote the initial manuscript. F.A. reviewed and edited the manuscript with contributions from C.C. All authors read and approved the final manuscript.

## Acknowledgements

We would like to thank Richard Stark (University of Warwick Bioinformatics & Digital Health RTP), Jack Weaver, and Tosin Orababa for their help with the installation and the operation of the statistical and genome assembly software packages. Lijiang Song is also acknowledged for help with UHPLC-HRMS analysis.

## Notes

### Competing Interest Statement

The authors have declared no competing interest.

