## Supplementary Information for "Identification of a terpene synthase arsenal using long-read sequencing and genome assembly of *Aspergillus wentii*"

Supplementary Table 4: NCBI BioProject ID submission and Accession number of A. wentii CBS 141173. . 6

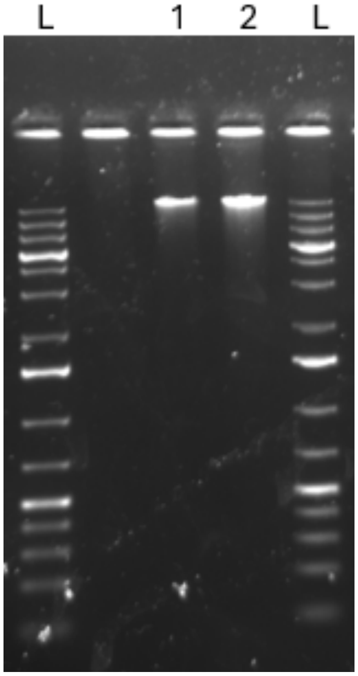

**Supplementary Figure 1: High Molecular Weight (HMW) gDNA from *A. wentii* CBS 141173**. Lanes L = 1kb plus ladder (Thermo Scientific). Lane 1 and 2= gDNA from *A. wentii*.

**Supplementary Table 1: Concentration of High Molecular Weight (HMW) *A. wentii* gDNA.**

| Sample | Before ethanol precipitation (ng/µL) | After ethanol precipitation (ng/µL) |
|---|---|---|
| *Aspergillus wentii* CBS 141173 - 1 | 65.4 | 481.2 |
| *Aspergillus wentii* CBS 141173 - 2 | 60.6 | 475.5 |

**Supplementary Table 2: Summary of RNA-seq analysis report of *Aspergillus wentii* CBS 141173**

| Sample ID | Raw PE Reads | Read pairs after deduplication and quality trimming | % Duplication Rate | Total Mapped Reads | & Total Mapped Reads | Unique Mapped Reads | % Unique Mapped Reads |
|---|---|---|---|---|---|---|---|
| **Awentii7A** | 14,193,289 | 12,162,221 | 12.99 | 9,407,395 | 77.35 | 9,338,336 | 76.78 |
| **Awentii7B** | 14,009,686 | 11,972,577 | 13.35 | 9,440,227 | 78.85 | 9,373,124 | 78.29 |
| **Awentii30A** | 12,697,660 | 10,745,915 | 14.07 | 8,623,884 | 80.25 | 8,575,636 | 79.80 |
| **Awentii30B** | 13,059,646 | 11,199,580 | 12.89 | 8,887,091 | 79.35 | 8,837,628 | 78.91 |

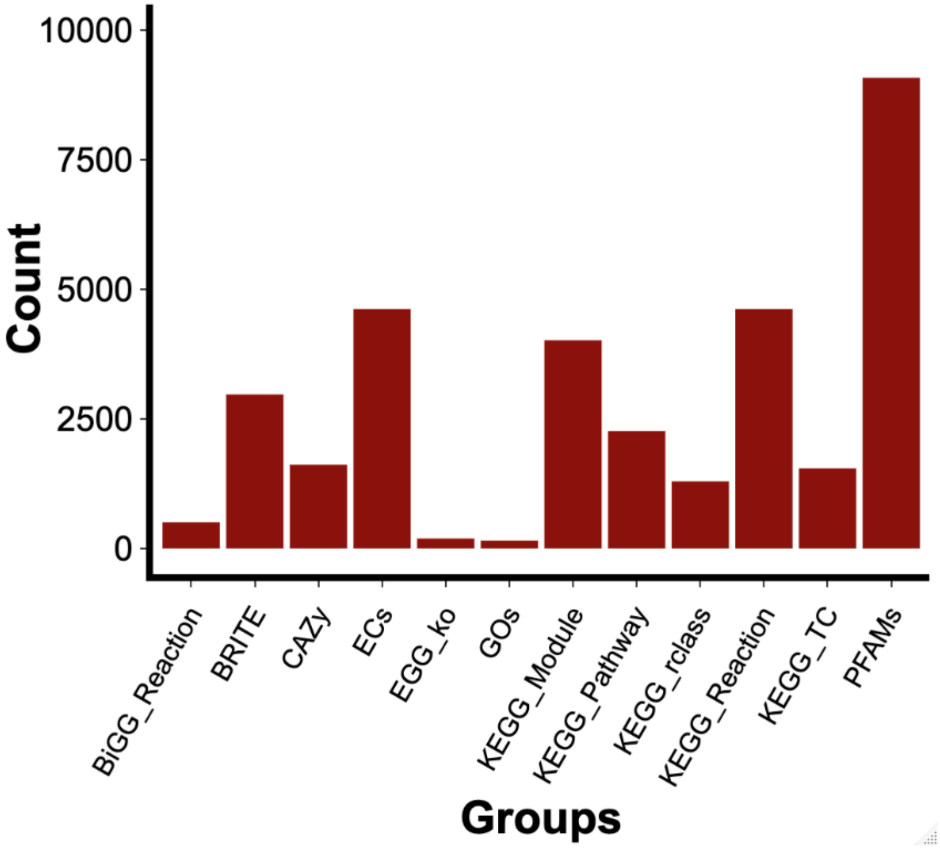

**Supplementary Figure 2: Assignment of predicted proteins in *A. wentii* CBS 141173 using eggnog mapper to multiple databases.**

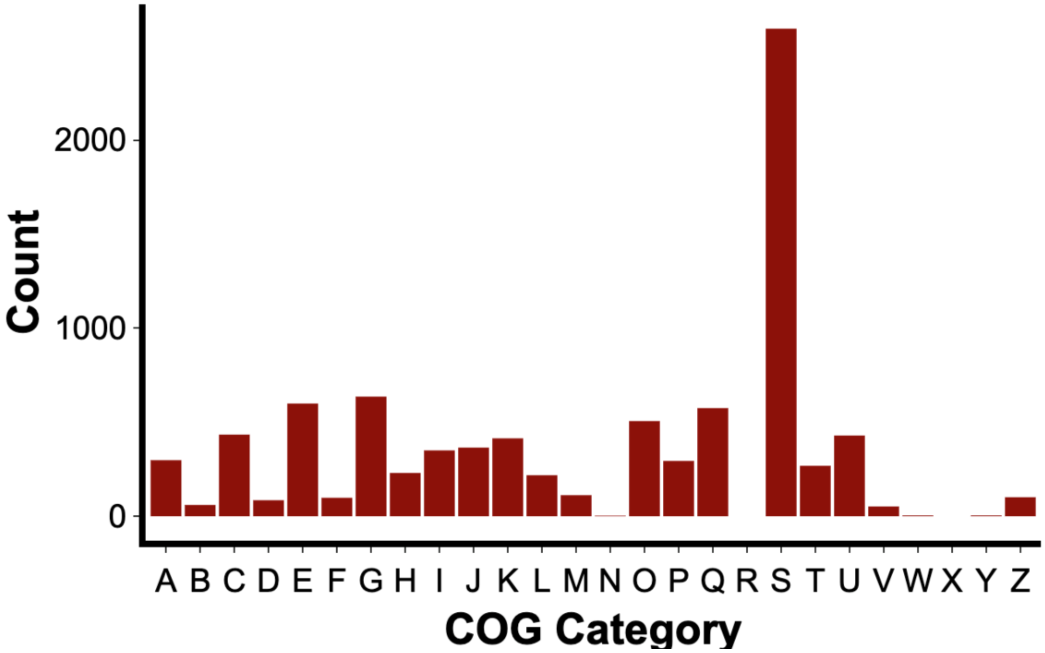

**Supplementary Figure 3. Distribution of predicted proteins in *Aspergillus wentii* CBS 141173 across the different COG categories.** A: RNA processing and modification. B: Chromatin structure and dynamics. C: Energy production and conversion. D: Cell cycle control, cell division, chromosome partitioning. E: Amino acid transport and metabolism. F: Nucleotide transport and metabolism. G: Carbohydrate transport and metabolism. H: Coenzyme transport and metabolism. I: Lipid transport and metabolism. J: Translation, ribosomal structure. and biogenesis. K: Transcription. L: Replication, recombination. and repair. M: Cell wall/membrane/envelope biogenesis. N: Cell motility. O: Posttranslational modification, protein turnover, chaperones. P: Inorganic ion transport and metabolism. Q: Secondary metabolites biosynthesis, transport. and catabolism. S: Function unknown. T: Signal transduction mechanisms. U: Intracellular trafficking, secretion, and vesicular transport. V: Defense mechanisms. W: Extracellular structures. Y: Nuclear structure. Z: Cytoskeleton. The following returned no hits: R: General function prediction only; X: Mobilome- prophages, transposons.

**Supplementary Table 3: BUSCO assessment of the annotated genome and predicted proteome and transcriptome of *A. wentii* CBS 141173 using the Eurotiales odb10 database.**

| Parameters | Genome | Protein | Transcriptome |
|---|---|---|---|
| **Percentage BUSCO** | 94.8% | 95.3% | 94.8% |
| **Complete BUSCO’s** | 3973 | 3992 | 3973 |
| **Complete and single copy BUSCO’s** | 3960 | 3751 | 3960 |
| **Complete and duplicated BUSCOs** | 13 | 241 | 13 |
| **Fragmented BUSCOs** | 27 | 120 | 27 |
| **Missing BUSCOs** | 191 | 79 | 191 |
| **Total BUSCO groups searched** | 4191 | 4191 | 4191 |

**Supplementary Table 4: NCBI BioProject ID submission and Accession number of *A. wentii* CBS 141173.**

| SUBID | BioProject | BioSample | Localid | Accession |
|---|---|---|---|---|
| **SUB14389936** | PRJNA1133277 | SAMN42381744 | scaffold_1 | CP165651 |
| **SUB14389936** | PRJNA1133277 | SAMN42381744 | scaffold_2 | CP165652 |
| **SUB14389936** | PRJNA1133277 | SAMN42381744 | scaffold_3 | CP165653 |
| **SUB14389936** | PRJNA1133277 | SAMN42381744 | scaffold_4 | CP165654 |
| **SUB14389936** | PRJNA1133277 | SAMN42381744 | scaffold_5 | CP165655 |
| **SUB14389936** | PRJNA1133277 | SAMN42381744 | scaffold_6 | CP165656 |
| **SUB14389936** | PRJNA1133277 | SAMN42381744 | scaffold_7 | CP165657 |
| **SUB14389936** | PRJNA1133277 | SAMN42381744 | scaffold_8 | CP165658 |
