## Supplementary Dataset 3 for "Identification of a terpene synthase arsenal using long-read sequencing and genome assembly of *Aspergillus wentii*"

**Terpenoid gene clusters from *Aspergillus wentii* CBS 141173**

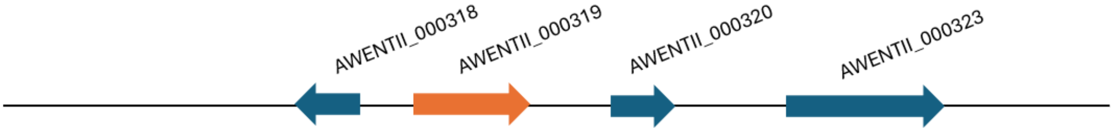


Table 1: AwTS1 terpenoid gene cluster

| **Gene locus tag** | **Homologue (% identity/% similarity), Organism, accession number** |
|---|---|
| AWENTII_000319 | Squalene hopane cyclase *afumA* (50/64) *Aspergillus fumigatus* A1163, B0Y565.1 |
| AWENTII_000318 | Cytochrome P450 monooxygenase afumB, (37/53) *Aspergillus fumigatus* A1163, B0Y566.1 |
| AWENTII_000320 | Acetyltransferase *adrJ,* Andrastin A biosynthesis cluster, (35/55), *Penicillium roqueforti*, A0A1Y0BRF4.1, |
| AWENTII_000323 | ABC multidrug transporter *mdr1*, (40/57) *Aspergillus fumigatus* Af293, Q4WTT9.1 |

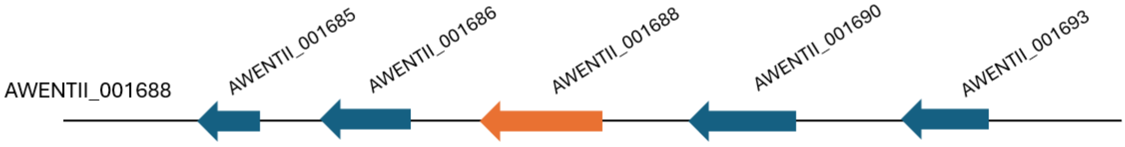


Table 2: AwTS2 terpenoid gene cluster

| **Gene Locus tag** | **Homologue (% identity/% similarity), Organism, accession number** |
|---|---|
| AWENTII_001688 | bicyclogermacrene synthase (31/45) *Penicillium expansum*, A0A0A2JP58.1 |
| AWENTII_001685 | Fatty acid hydroxylase *vlmA,* (44/60) *Lecanicillium sp,* A0A024FA41.1 |
| AWENTII_001686 | L-amino-acid oxidase, (38/55) *Neurospora crassa* OR74A, P23623.2 |
| AWENTII_001690 | D-threonine aldolase, (39/54), *Arthrobacter* sp. O82872.1 |
| AWENTII_001693 | Serine/threonine-protein phosphatase 6 regulatory ankyrin repeat subunit C (31/46), Q502K3.1 |

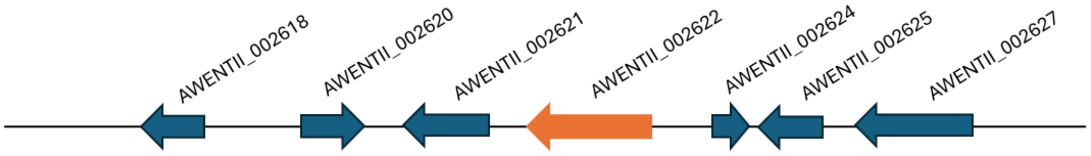


Table 3: AwTS3 terpenoid gene cluster

| **Gene Locus Tag** | **Homologue (% identity/% similarity), Organism, accession number** |
|---|---|
| AWENTII_002622 | Lanosterol synthase *erg*7A (89/92) *Aspergillus fumigatus* Af293, Q4WES9.1 |
| AWENTII_002618 | Serine/threonine-protein kinase PHO85 (77/86), *Candida albicans*, Q9HGY5.1 |
| AWENTII_002620 | Vacuolar histidine transporter YPQ3, (38/57), *Saccharomyces cerevisiae* S288C, P38279.1, |
| AWENTII_002621 | Glucose-insensitive transcription protein 7 (25/41), *Schizosaccharomyces pombe* 972h- O59709.2 |

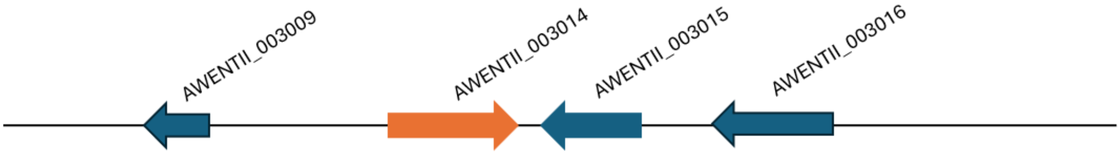


Table 4: AwTS4 terpenoid gene cluster

| **Gene locus tag** | **Homologue (% identity/% similarity), Organism, accession number** |
|---|---|
| AWENTII_003009 | Serine/threonine-protein phosphatase 6 regulatory ankyrin repeat subunit C |
| AWENTII_003014 | Terpene cyclase (24/42) *Fusarium fujikuroi* IMI 58289, S0EGZ9.1 |
| AWENTII_003015 | FAD-dependent oxidoreductase (46/65) *Aspergillus flavus* NRRL3357 |
| AWENTII_003016 | High-affinity glucose transporter (43/63), Schizosaccharomyces pombe 972h |

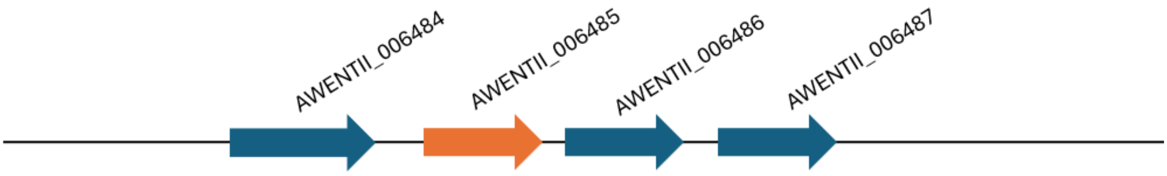


Table 5: AwTS5 terpenoid gene cluster

| **Gene locus** | **Homologue (% identity/% similarity), Organism, accession number** |
|---|---|
| AWENTII_006486 | Lanosterol synthase *erg7A* (41/59) *Aspergillus fumigatus* Af293, Q4WES9.1 |
| AWENTII_006484 | ubiquitin-protein ligase (43/56) *Mus musculus* Q6ZQ89.2 |
| AWENTII_006485 | ERAD-associated E3 ubiquitin-protein ligase doa10 (30/52) *Schizosaccharomyces pombe* 972h, O60103.1 |
| AWENTII_006487 | Probable inactive receptor kinase At3g02880 (25/37) *Arabidopsis thaliana,* Q9M8T0.1 |

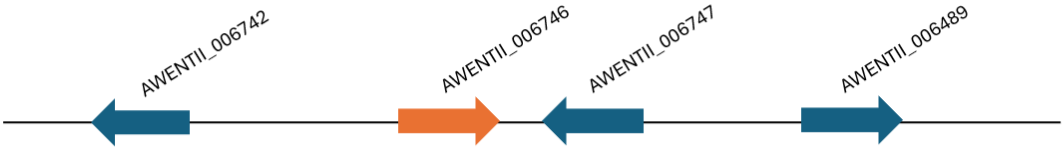


Table 6: AwT6 terpenoid gene cluster

| **Gene Locus** | **Homologue (% identity/% similarity), Organism, accession number** |
|---|---|
| AWENTII_006742 | Short-chain dehydrogenase ptmH (41/61) *Penicillium simplicissimum*, A0A140JWS5.1 |
| AWENTII_006746 | Bifunctional lycopene cyclase (59/66) *Nannizzia gypsea* CBS 118893, E4UPP6.1 |
| AWENTII_006747 | Phytoene desaturase (59/74), *Fusarium fujikuroi* IMI 58289, S0EPU6.1 |
| AWENTII_006749 | Short-chain dehydrogenase fogG, (33/51) *Aspergillus ruber* CBS 135680 A0A017SEY2.1 |

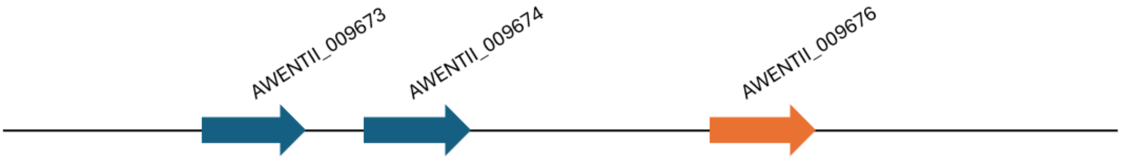


Table 7: AwTS7 terpenoid gene cluster

| **Gene locus** | **Homologue (% identity/% similarity), Organism, accession number** |
|---|---|
| AWENTII_009673 | Maleylacetate reductase (52/67), *Burkholderia cepacian*, Q45072.1 |
| AWENTII_009674 | S-adenosyl-L-methionine:L-histidine 3-amino-3-carboxypropyltransferase (80/89), *Aspergillus fumigatus* Af293, Q4WN99.1 |
| AWENTII_009676 | (E)-beta farnesene synthase MBR_03882 (32/52) *Metarhizium brunneum* ARSEF 3297, A0A0B4G504.1 |

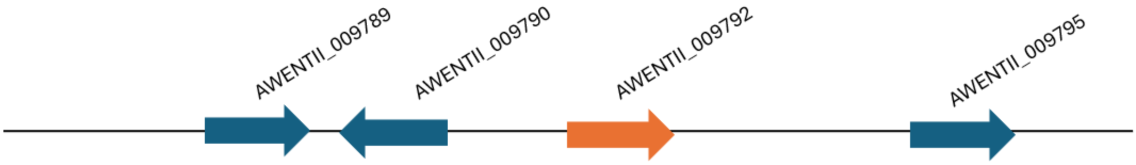


Table 8: AwTS8 terpenoid gene cluster

| **Gene locus** | **Homologue (% identity/% similarity), Organism, accession number** |
|---|---|
| AWENTII_009789 | Cytochrome P450 monooxygenase *drtD* (43/60) *Aspergillus calidoustus,* A0A0U5GRB4.1 |
| AWENTII_009790 | Cytochrome P450 monooxygenase *penB* (35/52), *Penicillium thymicola*, A0A1B2CTB6.1 |
| AWENTII_009792 | Squalene hopane cyclase afumA (44/58) *Aspergillus fumigatus* A1163, B0Y565.1 |
| AWENTII_009795 | Acetyltransferase FGR3 (63/77), *Fusarium graminearum* PH-1, I1RLA5.1 |

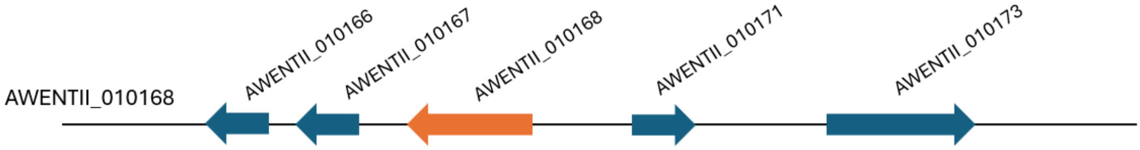


Table 9: AwTS9 terpenoid gene cluster

| **Gene locus** | **Homologue (% identity/% similarity), Organism, accession number** |
|---|---|
| AWENTII_010168 | Geranylgeranyl pyrophosphate synthase (66/82) Neurospora crassa OR74A, P24322.2 |
| AWENTII_010166 | G-protein complex alpha subunit gpaA (99/99),*Aspergillus fumigatus* A1163 B0XRA0.1 |
| AWENTII_010167 | Smr domain-containing protein (38/59) *Schizosaccharomyces pombe* 972h, Q9UTP4.1 |
| AWENTII_010171 | methyltransferase AN0656 (54/67) *Aspergillus nidulans* FGSC A4, Q5BFM4.1 |

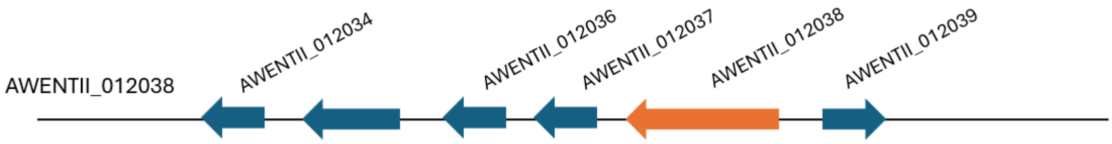


Table 10: AwTS10 terpenoid gene cluster

| **Gene locus** | **Homologue (% identity/% similarity), Organism, accession number** |
|---|---|
| AWENTII_012034 | 2-oxoglutarate dehydrogenase complex component E2 (50/65), *Schizosaccharomyces pombe* 972h O94681.1 |
| AWENTII_012036 | Acid protease A (43/58), *Aspergillus niger,* P24665.1 |
| AWENTII_012037 | Dimeric xanthone (28/44), *Cryptosporiopsis* sp. 8999, A0A4P8DJU7.1 |
| AWENTII_012038 | Farnesyl-diphosphate farnesyltransferase erg9 (79/88) *Aspergillus fumigatus* Af293, Q4WAG4.1 |
| AWENTII_012039 | Chaperone protein dnaJ (39/57), *Saccharomyces cerevisiae* S288C, P40564.1 |

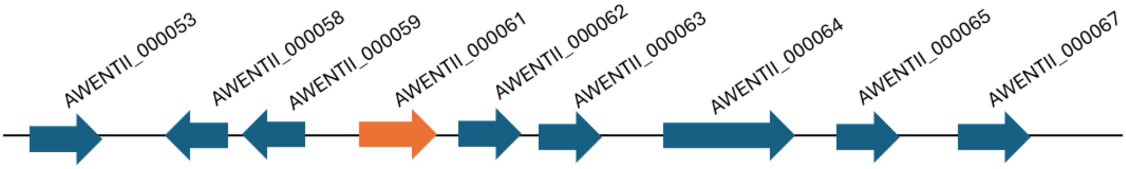


Table 11: AwTS11 terpenoid gene cluster

| **Gene ID** | **Homologue (% identity/% similarity), Organism, accession number** |
|---|---|
| AWENTII_000053 | 2-oxoglutarate-Fe(II) type oxidoreductase *ppzD* (28/42), *Metarhizium rileyi* RCEF 4871 |
| AWENTII_000058 | Thioesterase *poxG* (36/54), *Penicillium oxalicum* 114-2, S7ZEI0.1 |
| AWENTII_000059 | Cytochrome P450 monooxygenase a*drA* (53/69), *Penicillium rubens* Wisconsin 54-1255, B6HUQ4.1 |
| AWENTII_000061 | Terpene synthase *nvfL* (39/56), *Aspergillus novofumigatus* IBT 16806, A0A2I1BT01.1 |
| AWENTII_000062 | FAD-dependent monooxygenase *andE* (56/73), *Aspergillus stellatus*, A0A097ZPF7.1 |
| AWENTII_000063 | Prenytransferase *adrG* (50/69), *Penicillium roqueforti*, A0A1Y0BRF7.1 |
| AWENTII_000064 | Non-reducing polyketide synthase *andM* (43/61), *Aspergillus stellatus*, A0A097ZPE0.1 |
| AWENTII_000065 | O-methyltransferase *atr3* (40/54), *Stereocaulon alpinum*, A0A8F4PN06.1 |
| AWENTII_000067 | Vanillyl-alcohol oxidase (66/79), *Penicillium simplicissimum*, P56216.1 |

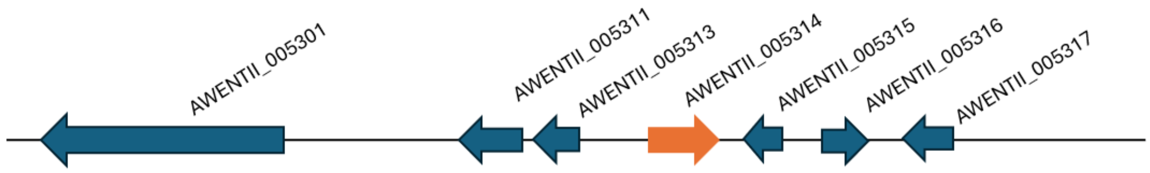


Table 12: AwTS12 terpenoid gene cluster

| **Gene locus** | **Homologue (% identity/% similarity), Organism, accession number** |
|---|---|
| AWENTII_005301 | Nonribosomal peptide synthetase *ungA* (38/55) *Aspergillus campestris* IBT 28561, A0A2I1D2N0.1 |
| AWENTII_005311 | Transcription factor FBD3 (24/38), *Fusarium pseudograminearum* CS3096, K3UIH8.1 |
| AWENTII_005313 | Cytochrome P450 monooxygenase *braC,* (34/51), *Annulohypoxylon truncatum,* P9WER2.1 |
| AWENTII_005314 | Fusicoccadiene synthase (29/47) *Diaporthe amygdali*, A2PZA5.1 |
| AWENTII_005315 | Xylitol dehydrogenase A (53/69), *Aspergillus fischeri* NRRL 181, A1D9C9.1 |
| AWENTII_005316 | L-arabinitol 4-dehydrogenase (51/64), *Penicillium rubens* Wisconsin 54-1255, B6HI95.1 |
| AWENTII_005317 | Dehydrogenase OXI1 (45/60), *Bipolaris maydis* ATCC 48331, N4WE73.1 |

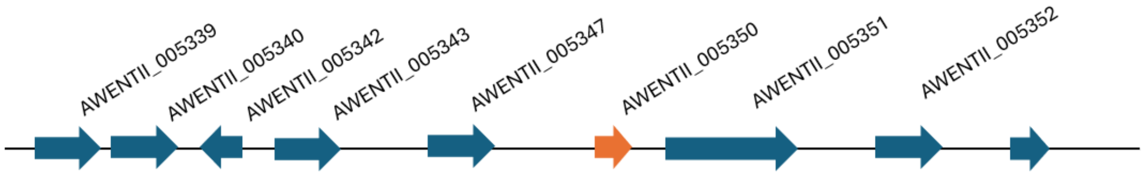


Table 13: AwTS13 terpenoid gene cluster

| **Gene locus** | **Homologue (% identity/% similarity), Organism, accession number** |
|---|---|
| AWENTII_005339 | Oxidase ucsJ (30/49), *Acremonium* sp. A0A411KUU5.1 |
| AWENTII_005340 | Flavin oxidoreductase hxnT (42/59), *Aspergillus nidulans* FGSC A4, A0A1U8QTA2.1 |
| AWENTII_005342 | Dehydrogenase (32/46), *Armillaria gallica*, A0A2H3D1U1.1 |
| AWENTII_005344 | FAD-dependent monooxygenase atA (54/70), *Aspergillus terreus* NIH2624, Q0CJ62.1 |
| AWENTII_005345 | Inositol 2-dehydrogenase/oxidoreductase (29/50), *Bacillus subtilis* subsp. subtilis str. 168, P40332.2 |
| AWENTII_005346 | Transcription factor atnE, (21/34) *Arthrinium* sp. A0A455M2Z1.1 |
| AWENTII_005347 | Efflux pump atB (72/82), *Aspergillus terreus* NIH2624, Q0CJ61.1 |
| AWENTII_005350 | Terpene cyclase flvE, (34/50) *Aspergillus flavus* NRRL3357, B8NHE0.1 |
| AWENTII_005351 | Non-canonical non-ribosomal peptide synthetase (29/49), *Fusarium verticillioides* 7600, W7N2C1.1 |
| AWENTII_005353 | Alpha-acetolactate decarboxylase, (32/53), *Bacillus* subtilis subsp. subtilis str. 168, Q04777.1 |

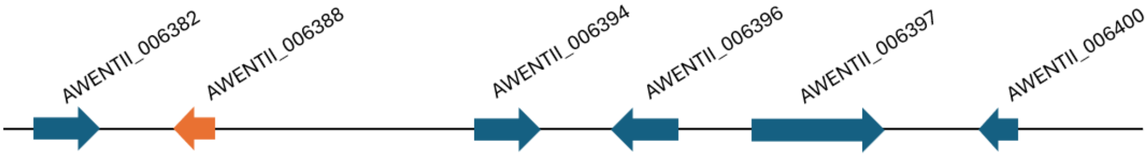


Table 14: AwTS14 terpenoid cluster

| **Gene Locus** | **Homologue (% identity/% similarity), Organism, accession number** |
|---|---|
| AWENTII_006382 | Oxygen-dependent choline dehydrogenase (28/44), *Yersinia pseudotuberculosis*, B2K8U4.1 |
| AWENTII_006388 | (E)-beta farnesene synthase MBR_03882 (35/54), Metarhizium brunneum ARSEF 3297, A0A0B4G504.1 |
| AWENTII_006394 | Methyltransferase cfoC (32/45), *Aspergillus candidus*, A0A2I2F2K7.1 |
| AWENTII_006396 | Cytochrome P450 monooxygenase xanG (41/62), *Aspergillus fumigatus* Af293, Q4WED5.1 |
| AWENTII_006397 | Isocyanide synthase xanB (63/79), *Aspergillus fumigatus* Af293, Q4WED9.2 |
| AWENTII_006400 | Phosphoglycolate phosphatase (29/45), *Pectobacterium atrosepticum* SCRI1043, Q6CZR3.1 |

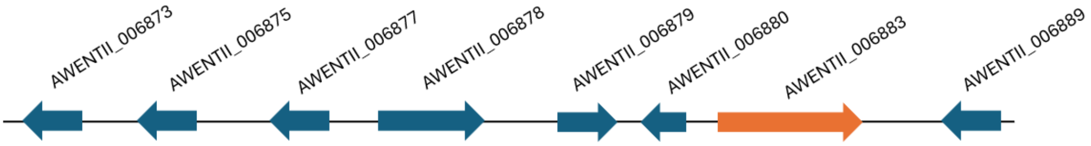


Table 15: AwTS15 terpenoid gene cluster

| **Gene locus** | **Homologue (% identity/% similarity), Organism, accession number** |
|---|---|
| AWENTII_006873 | Hydroxymethylglutaryl-CoA synthase, (60/75) *Aspergillus fumigatus* Af293, Q4WXT8.1 |
| AWENTII_006875 | 3-hydroxy-3-methylglutaryl coenzyme A reductase, (78/89), *Aspergillus nidulans* FGSC A4, C8VN86.1 |
| AWENTII_006876 | Transcription factor pbcR (54/66), *Aspergillus nidulans* FGSC A4, A0A1U8QL22.1 |
| AWENTII_006877 | Cytochrome P450 monooxygenase (82/88), *Aspergillus nidulans* FGSC A4, C8VN91.1 |
| AWENTII_006879 | Oxidoreductase (59/77), *Aspergillus nidulans* FGSC A4, A0A1U8QJR1.1 |
| AWENTII_006880 | Oxidoreductase (80/90), *Aspergillus nidulans* FGSC A4, A0A1U8QP15.1 |
| AWENTII_006883 | Pimaradiene synthase (61/75), *Aspergillus nidulans* FGSC A4, A0A1U8QHE3.1 |
| AWENTII_006889 | Uncharacterised protein |

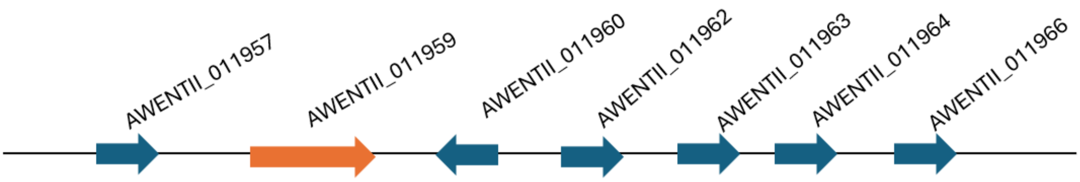


Table 16: AwTS16 terpenoid gene cluster

| **Gene Locus** | **Homologue (% identity/% similarity), Organism, accession number** |
|---|---|
| AWENTII_011957 | Cytochrome P450 monooxygenase olcB (43/58), *Penicillium canescens*, P9WEQ1.1 |
| AWENTII_011959 | Pimaradiene synthase pbcA (43/60), *Aspergillus nidulans* FGSC A4, A0A1U8QHE3.1 |
| AWENTII_011960 | Lactonohydrolase oryL (30/46), Aspergillus oryzae RIB40 Q2TXF9.2 |
| AWENTII_011962 | Cytochrome P450 monooxygenase tpeC (41/60), *Talaromyces stipitatus* ATCC 10500, B8MV61.1 |
| AWENTII_011963 | Transcription factor pbcR (26/44), *Aspergillus nidulans* FGSC A4, A0A1U8QL22.1 |
| AWENTII_011964 | Cytochrome P450 monooxygenase ntnM (39/55), *Fusarium fujikuroi* IMI 58289, S0E2U7.1 |
| AWENTII_011966 | 4-hydroxyproline 2-epimerase (44/63), *Brucella anthropi* ATCC 49188, A6WW16.1 |
